# DNA origami assembled spheroid for evaluating cytotoxicity and infiltration of chimeric antigen receptor macrophage (CAR-M)

**DOI:** 10.1101/2023.12.03.569750

**Authors:** Junhua Liu, Xiaofang He, Qingyao Zhu, Heming Wang, Xiaojiao Shan, Yicheng Zhao, Luo Zhang, Guangqi Song, Xiushan Yin

**Author notes:** Corresponding author. NHC Key Laboratory of Biotechnology Drugs (Shandong Academy of Medical Sciences), Biomedical Sciences College, Shandong First Medical University, Ji’nan 250117, China. Corresponding author. PuHeng Biotechnology (Suzhou) Co., Ltd. 215000, China. Corresponding author. Research Center of Bioengineering, the Medical Innovation Research Division of Chinese PLA General Hospital, Beijing 100039, China. E-mail addresses:* (Xiushan Yin); (Guangqi Song) and (Luo Zhang). These authors contributed to this work equally as co-authors (Junhua Liu, Xiaofang He, Qingyao Zhu, Heming Wang).

## Abstract

Chimeric antigen receptor (CAR) T-cell therapies have shown remarkable results in patients with hematological malignancies. However, their success in treating solid tumors has been limited. As an alternative candidate for the CAR approach, CAR-macrophages (CAR-M) have demonstrated activation and phagocytosis directed by tumor antigens, showing promise in the treatment of solid tumors. Nevertheless, the mechanisms by which CARs direct tumor chemotaxis and invasion of CAR-M remain poorly understood. In this study, we aim to investigate the role of CARs in CAR-M attachment and infiltration using 3D tumor micro-spheroids, which were created by utilizing a novel nucleic acid nanostructures decorated living cells (NACs) based origami assembly technique. First, the effectiveness of phagocytosis and killing conducted by CAR-M was validated in the conventional 2D well/plate-surface culture models. Then,peripheral blood mononuclear cell (PBMC) invasion assay confirmed that the 3D tumor micro-spheroids were feasible for cell invasion. Finally, our results demonstrated that CAR-M exhibited higher invasion and killing capacity in 3D tumor micro-spheroids. In summary, the 3D NACs-origami assembled tumor spheroid model provides a suitable platform for target screening and pharmacodynamic evaluation of CAR-M.

## 1. Introduction

In recent years, immunotherapy has revolutionized cancer treatment. CAR-T, as the first personalized cellular therapeutic strategy, has successfully been used to treat refractory hematologic malignancies, particularly those targeting CD19 [1, 2]. However, there are obstacles in treating solid tumors, such as low infiltration and high heterogeneity in the tumor microenvironments (TME) [3], To address this, CAR-NK (Natural Killer) cell therapy has emerged as a promising alternative strategy, demonstrating efficacy in targeting solid tumor antigens in clinical trials [4-6]. Another alternative strategy is CAR-M (CAR macrophage) therapy, which has shown advantages in immune cell trafficking and infiltration effect [7, 8]. For example, Michael Klichinsky and colleagues delivered CAR structures into human primary macrophages using adenovirus transduction, resulting in a unique phagocytic effect, enhanced expression of proinflammatory factors, and cross-presentation of tumor antigens to bystander T cells in vivo [9]. Other studies have focused on enhancing phagocytosis to improve the anti-tumor activity of CAR-macrophages [10-12]. This includes combining anti-CD47 antibody to affect CD47-Sirpα axis interaction and enhance phagocytosis [11], optimizing the intracellular PI3K-signaling/Megf7 domain [11] and MerTK [12], and using exogenous chemicals to stimulate CAR-M towards an M1 pro-inflammatory phenotype, which enhances phagocytosis, tumor killing ability, and cytokine release on target cells [13].. In summary, CAR macrophages show promise as a potential treatment for solid tumors.

Animal tumor models are commonly used for evaluating the efficacy and safety of cell therapies in a preclinical setting. However, these models are time-consuming, expensive, and raise ethical concerns. As an alternative, 3D tumor spheroids that simulate the tumor microenvironment in vitro have emerged as a promising model for preclinical tumor treatment studies. Extensive research evidence suggests that 3D tumor spheroid/organoid models can serve as effective methods for preclinical drug screening and verification [14, 15]. Various 3D tumor spheroid/organoid models have been reported for different solid tumors, including pancreatic cancer [16], ovarian cancer [17, 18], colon cancer [19], liver cancer [20], prostate cancer [21], and bladder cancer [22]. However, there are limited reports on the use of 3D tumor spheroid models to evaluate CAR-mediated immune cell therapy [23-25]. One study demonstrated the co-culture of CAR-T cells with spheroids from a colorectal cancer cell line to mimic the complex morphology of solid tumors, but this approach was limited to a specific type of adherent cell line [23]. To address these challenges, we propose a novel approach using DNA origami techniques to assemble a 3D tumor spheroid system. DNA origami is based on the pairing principle of DNA strands, where a desired DNA single-long strand, known as the ‘scaffold strand’, is folded into a designed cluster of oligonucleotide strands called ‘staple strands’ [26-28]. This allows nanoscale DNA origami to self-assemble on target cells and facilitate interactions among multiple target cells. Building upon our previously published work on 3D nucleic acid nanostructures to decorate living cells (NACs) using origami assembly techniques to form 3D liver spheroids [29, 30], we aim to investigate the role of CARs in CAR-M attachment and infiltration using 3D tumor micro-spheroids based on origami assembly technique.

In this study, we used a unique NACs-based origami assembly technique to model 3D tumor micro-spheroids. We evaluated the biological functional characteristics of CAR-M not only in 2D well/plate-surface cultures but also in a 3D tumor spheroid model. These experimental data confirmed the targeted recruitment, infiltration, and killing effect of CAR-M on tumor spheroids in a complex and organized tumor microenvironment. In conclusion, the 3D NACs-origami assembled tumor spheroid model is well-suited for evaluating pharmaceutical efficacy and targets screening for CAR-M.

## 2. Materials and Methods

### 2.1. Cell Culture

RAW264.7, ovarian cancer cell SKOV3 and 293FT cells were purchased from ATCC. RAW264.7 (also known as RAW), RAW264.7 GFP, and RAW264.7 CAR-HER2 GFP cells were cultured in RPMI-1640 medium (Corning), supplemented with 10% fetal bovine serum (Lonza) and 100 U/ml penicillin/streptomycin (Life Technologies). 293FT, SKOV3 and SKOV3-fluc cells were cultured in Dulbecco’s Modified Eagle Medium(Corning), also supplemented with 10% FBS and 100 U/ml penicillin/streptomycin. Above all cells were cultured in 5% CO2 incubators at 37 □. For the subsequent cell analysis experiment, RAW264.7 cells were collected by direct pipetting, and the other adherent cells were digested by 0.25% Trypsin-EDTA (Gibco).

The mouse peripheral blood mononuclear cells (PBMCs) in our study were purchased from Oricells company, and finished product was namely mouse PB-MNC (MOSPB-001F). They were cultured in a complete RPMI medium to ensure their vitality.

### 2.2 Lentivirus production and infection

Preparation of CAR-M cells and SKOV3-fluc tumor cells were transducted by CAR lentivirus particles and luciferase lentivirus particles, respectively. The lentiviruses were produced by transfecting 293FT cells with a thirdgeneration plasmid system, including pMD2.G, psPAX2, and a lentiviral expression vector. The 293FT cells were seeded and transfected using the jetPRIME transfection reagent (Polyplus). 48 hours and 72 hours post-transfection, cell supernatants were collected and centrifuged at 250 g for 5 minutes to remove cell debris. The above lentivirus supernatant was then filtered through a 0.45 μm PVDF filter and concentrated using LentiX (Takara Biosciences). The final lentivirus can be added directly to the cells or stored in a -80 □ refrigerator.

### 2.3. Flow cytometry analysis

Multi-parameter flow cytometry (BD FACS catoII) was utilized to analyze CAR expression or collect other fluorescence signals. For the detection of CAR expression in CAR-M cells, CAR-M cells was prepared and the Fc receptor was blocked with anti-moCD16/32 antibody (BioLegend) for 10 minutes at room temperature. Subsequently, the cells were incubated with Myc tag-APC antibody(BioLegend) on ice for 30 minutes in the dark. After washing with the FACS buffer, the cells were measured on the flow cytometry machine. For the analysis of phagocytic efficiency, CellTracker DeepRed (Thermo Fisher Scientific), a fluorescent dye for labeling living cells, was used to label tumor cells at a working concentration of 1 μM per 10^6 cells/ml. Finally, the FCS data file exported by BD Diva software was further analyzed using FlowJo software(version 10.8.1).

### 2.4. Western blot analysis(WB)

Cell pellets were lysed by RIPA buffer containing PMSF on ice. The total protein concentration was quantified by a BCA kit (Beyotime), The lysates were then mixed with 10x loading buffer and denatured by boiling water. Each protein sample was separated on SDS-PAGE and transferred to PVDF membranes. The membrane was blocked by 5% skim milk in TBST buffer, and followed by incubation overnight at 4 □ with primary antibodies, anti-mouse Myc tag antibodies (1:1000, CST) or anti-β-actin monoclonal antibodies (1:3000, CST). On the second day, the membranes were washed three times and incubated with anti-mouse secondary antibodies (1:1000, CST) for 1 hour at room temperature. After washing, the membranes were developed with chemistry high-sig ECL Western Blotting substrate(Tanon) and analyzed with the chemiDoc XR (BIO-RAD).

### 2.5. Live Cell Staining

Cells were collected according to the routine procedures and centrifuged at 250 g for 5 minutes. The final cell pellet was resuspended in PBS or serum-free culture medium. Next, CellTracker Deep Red Dye (Thermo Fisher Scientific, C34565) or CellTracker Green CMFDA Dye (Thermo Fisher Scientific, C2925) was added to the suspension solution at a concentration of 1 μM per 10^6 cells/ml. The cell-dye mixtures were incubated at 37 °C in the dark. After staining for 30 minutes, the cells were washed twice with culture medium and can be used for co-culture experiments with CAR-M cells.

### 2.6. Phagocytosis Assay

Due to the intrinsic GFP fluorescent protein label in CAR-M cells, a live cell staining method was used to differentiate effector cells from target cells. Tumor cells were stained with CellTracker DeepRed (Thermo Fisher Scientific, C34565). Approximately 1 x 10^5 RAW264.7 CAR-HER2 GFP or RAW264.7 GFP cells were pre-seeded in a 6-well plate. Subsequently, an appropriate quantity of stained tumor cells was added in the dark to attain an effector to target(E/T) ratio of 1:3. After co-incubation for 3-6 hours, fluorescence microscopy imaging and flow cytometry analysis were performed respectively. Ultimately, the phagocytic efficiency of CAR-M cells was assessed by double-positive events of GFP and RED.

### 2.7. Tumor Killing Assay

Based on the Flyfire Luciferase fluorescence reporting system for evaluating killing efficiency, RAW264.7 CAR-HER2 GFP or RAW264.7 GFP were pre-seeded into a 96-well black plate (Beyotime) at a density of 10^4 cells/100 μl medium per well. Then target cells were added at different E/T ratios(100 μl/well) and co-cultured at 37□ in CO2 incubator for 24 hours or 48 hours. After incubation, each well was replaced with 100 μl cell medium containing D-Luciferin (substrate) at a working concentration recommended as 0.3 mg/ml. Next, the 96-well plates were incubated at 37 □for 5-10 minutes without light, and then transferred to a universal microplate reader for bioluminescence detection(Cytation1, Agilent). Based on the LDH method for evaluating killing efficiency, the cell cytotoxicity was converted by the formula as follows:

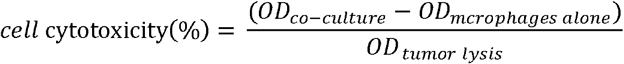

### 2.8. Construction of 3D tumor spheroid model

During the construction of the 3D tumor spheroid model, the 3D NACs-origami assembly technique was employed to replicate the three-dimensional growth environment characteristic of tumors. Initially, SKOV3-Fluc cells or SKOV3-mcherry cells were prepared. These cells were isolated from the culture dishes using enzymatic digestion. SKOV3-Fluc cells (3000 cells per well) were seeded into a 96-well plate. Subsequently, the plate was placed in 5% CO2 incubators at 37 □and incubated for 24 hours. During this incubation period, the cells aggregated and initiated the formation of three-dimensional structures, closely mimicking the growth environment of tumors.

### 2.9. Construction of co-cultivation model

Utilizing the NACs-origami assembly technique, a 3D model was meticulously constructed. PBMCs were collected, serving as the source of immune cells essential for the establishment of an immune response component. These PBMCs were subsequently introduced into the 3D tumor model culture medium to ensure their co-existence with tumor cells. Simultaneously, immune-modulating agents, namely the immune suppressor BLZ945 (MCE, HY-12768) and the immune recruiter SEW2871 (MCE, HY-W008947) were incorporated to emulate an immunomodulatory environment [31, 32]. Subsequently, the co-cultured 3D model was placed in a controlled cell culture incubator and incubated for 24 hours. During this incubation period, regular monitoring and recording of PBMCs’ invasive capability were conducted, alongside an assessment of the potential influence of immune-modulating agents. These observations served to evaluate the intricate cellular behaviors and interactions within the model system.

### 2.10. LDH assay

A volume of 100 μL of supernatant was carefully collected and placed into a transparent 96-well plate, ensuring uniform distribution across all wells. Subsequently, the LDH content in the supernatant was assayed using an LDH assay kit (CK12, provided by Tongren Chemical). Following this, absorbance readings were obtained at 490 nm using a spectrophotometer. The LDH’s relative concentration was determined by measuring the absorbance values and subsequently calculating the LDH concentration based on the standard curve provided by the assay kit.

### 2.11. ATP assay

The requisite number of tumor spheroid samples was aspirated and carefully placed into a clear 96-well plate. Subsequently, an equivalent volume of ATP assay reagent (Vazyme, DD1201) was added and thoroughly mixed with the samples. Following the addition of the assay reagent, the plate was placed on a shaker and agitated at 600 rpm under subdued lighting conditions for a duration of 30 minutes. After the incubation, the luminescence intensity of each well was measured using a microplate reader at the appropriate wavelength. Given that the generated chemical luminescence is directly proportional to ATP concentration, the ATP content in each sample was determined by measuring the luminescence intensity.

### 2.12. Histopathology

Tumor spheroid samples from the 3D model were meticulously collected, followed by overnight fixation in 4°C paraformaldehyde (PFA). Subsequently, a dehydration and embedding procedure was conducted. The samples were sequentially immersed in varying concentrations of alcohol to gradually remove moisture. Following this, the tumor spheroid samples were embedded in paraffin to ensure structural fixation and preservation of integrity. In preparation for subsequent analysis, 4 μm continuous sections were meticulously obtained using a microtome. Hematoxylin and eosin (H&E) staining was employed for observing the overall spheroid structure, DAPI staining was used to delineate cellular nuclei, and immunofluorescence staining was performed for the detection of specific proteins or antigens.

## 3. Result

### 3.1. Validation of CAR-M in 2D surface culture

To evaluate the phagocytic and killing ability of CAR-M in vitro in 2D well/plate-surface culture, we firstly designed a CAR construct, which contains an extracellular Myc tag and anti-HER2 scFv domain, a hinged CD8 region transmembrane domain and an intracellular CD3 costimulatory domain(Fig. 1A). Additionally, a GFP fluorescent label was also expressed along with a CAR unit. Next, we successfully transduced CAR construct using the lentiviral packaging system into the macrophage cell line RAW264.7 (referred to as RAW264.7 CAR-HER2 GFP). RAW264.7 wildtype cells and RAW264.7 GFP cells served as the control groups (Fig. 1B). Flow cytometry analysis further confirmed CAR expression on the RAW CAR-HER2 cells (Fig. 1C).

**Fig. 1.**
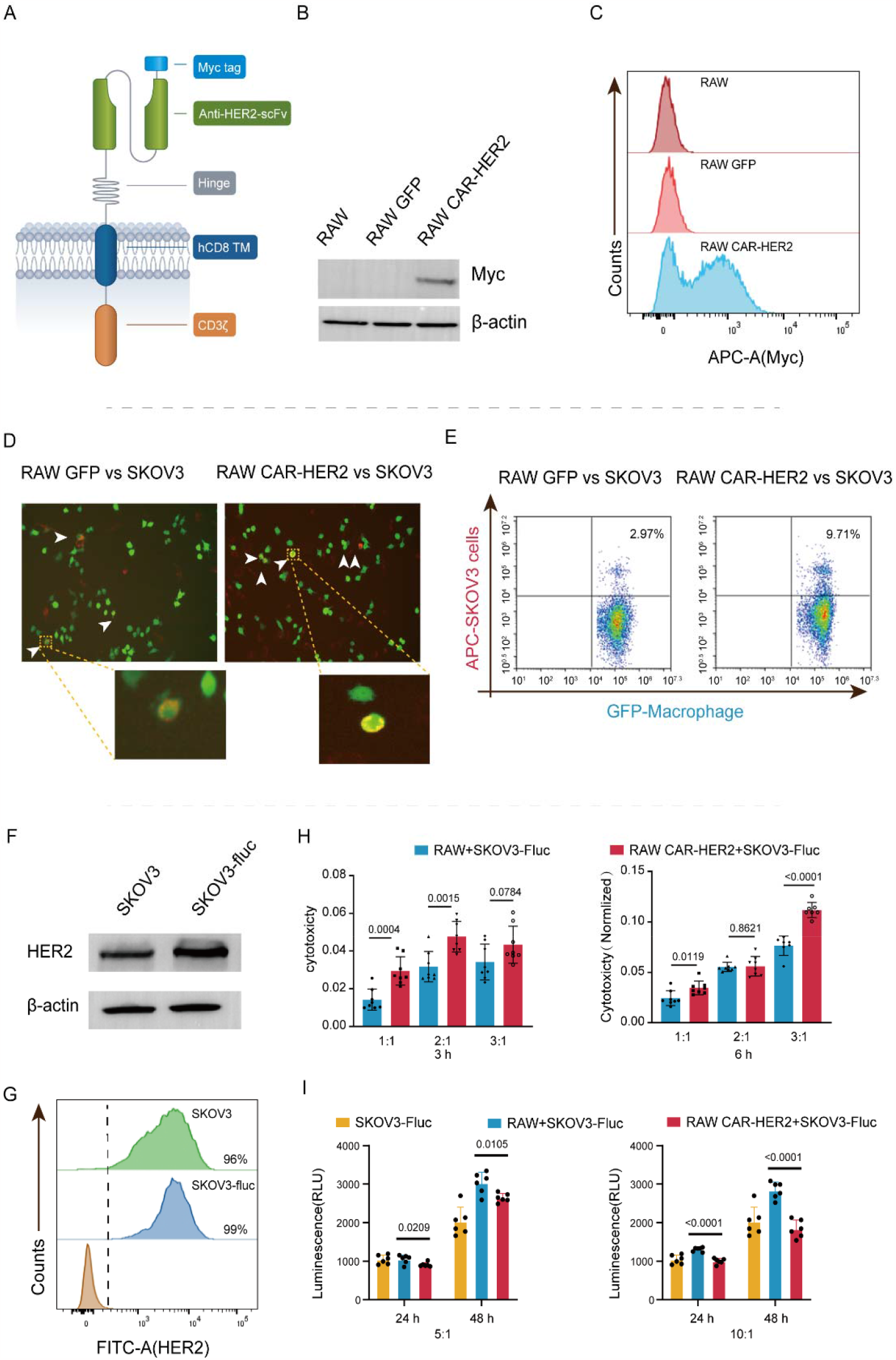
CAR-M establishment and 2D horizontal function verification. (1A) Schematic diagram of the CAR structure composition. (1B-C) Western blot analysis and flow cytometry analysis were used to show the CAR expression levels of RAW cells, RAW GFP cells, and RAW CAR-HER2 cells, respectively CAR structure includes a Myc tag. (1D) The phagocytic efficiency of RAW CAR-HER2 cells (or RAW GFP cells as a control group) against tumor cells SKOV3 (labeled with pHrodoSE) was demonstrated by microscopic imaging(x10 lens) after co-culture for 3 hours. Scale bar, 100 μm. (1E) The phagocytic efficiency of RAW CAR-HER2 cells (RAW GFP cells as a negative control group) against tumor cells SKOV3 (labeled with cell tracker FarRed) was demonstrated by flow cytometry analysis after co-culture for 6 hours. Efficiency percentage (%) refers to the macrophages counts occurred phagocytosis in all macrophages. (1F-G) Verification of the surface antigen HER2 expression in SKOV3 and SKOV3 Fluc cells via western blot analysis and flow cytometry analysis. (1H-I) Validation of the cytotoxic effect of RAW CAR-HER2 cells and RAW cells on targeting SKOV3 Fluc cells at different time points and different effector to targer (E/T) ratio via the LDH-based and firefly luciferase-based methods, respectively. P-values for statistical significance were determined by the t-test method(*P<0.05, **P<0.01, ***P<0.001, ****P<0.0001), and data are represented as the mean± SD of the n=6 technical replicates experiment.

Regarding the HER2 target, the ovarian cancer cell line SKOV3 was chosen for conducting functional experiments of CAR-M in vitro. To assess tumor-killing function, SKOV3 fluc cells with a luciferase reporter gene were utilized, which highly expressed HER2 (Fig. 1F, G) . In the pHrodo-based phagocytosis assay, SKOV3 tumor cells were labeled with pHrodoSE dye, which exhibits a red fluorescent signal upon phagocytosis and endocytosis occurring. Consequently, Our results suggested that after co-culturing CAR-M with SKOV3 tumor cells, a higher number of phagocytic events were observed in the RAW CAR-HER2 cells group than the control group (RAW264.7 GFP) in random picked fields of view via fluorescent microscope (Fig. 1D). In another phagocytosis assay using cell tracker FarRed Dye, flow cytometry analysis demonstrated that RAW CAR-HER2 cells showed high phagocytic efficiency (about 10%), which was superior to the negative control RAW264.7 GFP (about 2%) when RAW CAR-HER2 cells or RAW GFP were co-cultured with SKOV3 tumor cells labeled with cell tracker FarRed dye at a ratio of effector to target(E/T) (1:3) for 6 hours(Fig. 1E). Furthermore, to evaluate the killing ability of CAR-M against SKOV3 cells, two methods were employed: the LDH-based method (Fig. 1H) and the Firefly luciferase-based method (Fig. 1I), and both of which revealed significant killing ability of CAR-M. Meanwhile, the killing efficiency of CAR-M possessed a time-dependent tendency and was correlated to the ratio of E/T(Fig. 1H, I). In conclusion, CAR-M showed significant phagocytic and killing effects in the 2D well/plate-surface culture conditions.

### 3.2. Interaction between PBMC and 3D tumor micro-spheroids forms a tumor immune microenvironment

To replicate the invasion characteristics of tumor tissue more accurately by CAR-M cells, we utilized NACs-origami assembly technology to construct a 3D tumor immune microenvironment model (Fig. 2A). Considering the vital importance of cell activity in the success of experiments in the tumor model, we used PI staining to label dead cells and calcein-AM staining to label live cells (Fig. 2B), which revealed that there were almost no dead cells in the tumor model. Furthermore, we monitored the activity of tumor cells over a span of 5 days (Fig. 2C) and observed that ATP levels gradually increased over time, indicating that our tumor model possessed stable cell activity and continued to grow. To facilitate efficient CAR-M screening, we built an ovarian cancer tumor immune microenvironment model on a 384-well plate. Within just 24 hours, we observed a significant invasion of PBMC into the solid tumor spheroids (Fig. 2D). It is worth noting that our model also demonstrated that the invasion capability of PBMC into the tumor model can be regulated by immunomodulatory drugs. BLZ945, a CSF-1R inhibitor, reduced immune cell invasion into the tumor, while SEW2871, a S1P1 agonist, enhanced immune cell recruitment. After BLZ945 treatment, PBMC invasion into the model significantly decreased compared to the control group, whereas the SEW2871 treatment group showed a substantial increase. In conclusion, through the 3D tumor model we constructed, we successfully simulated the interaction between tumor and immune cells, allowing immune cells to actively invade the tumor model, mimicking the tumor immune microenvironment.

**Fig. 2.**
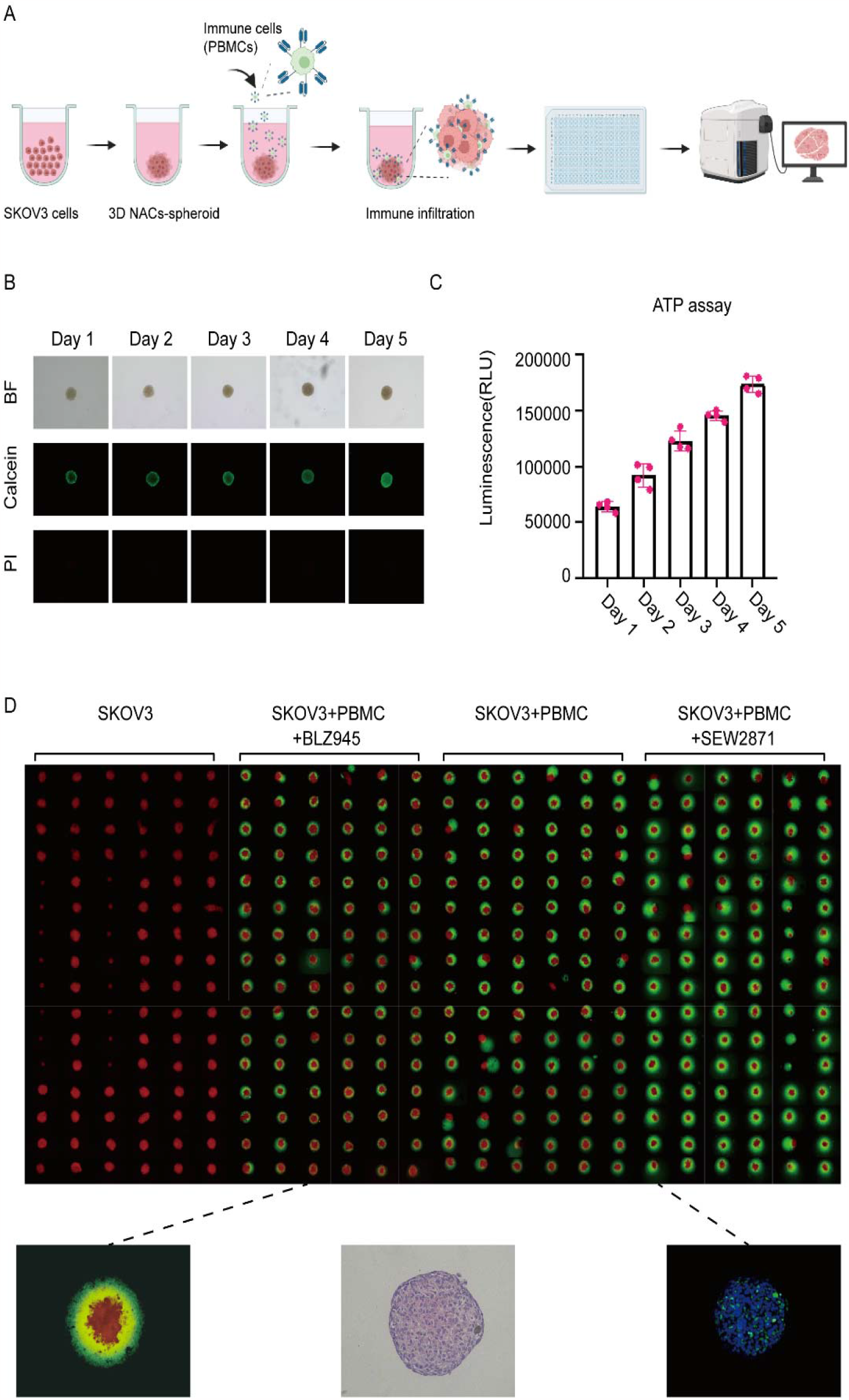
Interaction between PBMC and 3D NACs-origami assembled solid tumor spheroid. (2A)Schematic representation of the construction of a tumor immune model using 3D NACs-origami assembly technique. (2B) Calcein-AM/PI double staining of 3D ovarian tumor micro-spheroids of SKOV3. PI staining was used for dead cells, and Calcein-AM staining was used for live cells using x4 lens. Scale bar, 100 μm. BF indicates the bright-field. (2C) ATP level detection to assess the activity of the tumor model at different time points. Data are represented as the mean ±SD of n=4 technical replicates. (2D) High-throughput modeling in a 384-well plate. After 24 hours of model construction, mouse PBMCs were added, as well as a mixture of PBMCs with the immune cell inhibitor BLZ945, and a mixture of PBMCs with the immune stimulant SEW2871, to observe the invasion behavior of PBMCs into the solid tumor spheroid. Three images in the last row were selected from the PBMC group, including fluorescence imaging (x10), H&E staining (x20) and immunofluorescent staining for DAPI (x10). Scale bar, 100 μm.

### 3.3. Validation of CAR-M targeting, infiltration, and cytotoxicity in 3D micro-spheroids model

To validate the recognition, penetration, and cytotoxicity of CAR-HER2 macrophages against solid tumors, we established a solid tumor model using breast cancer SKOV3-Fluc cells. After successful model construction, we co-cultured RAW264.7-GFP and RAW264.7-CAR-HER2-GFP with the 3D NACs-origami assembled tumor spheroid (Fig. 3A). To determine the optimal ratio and time for interaction between tumor spheroid and RAW264.7 CAR-HER2 cells, we evaluated the infiltration and cytotoxic effects of macrophages at different ratios of tumor cells to macrophages(Fig. 3B). CAR-HER2 macrophages at various ratios could effectively infiltrate the tumor spheroid, with the E/T ratio of 10:1showing the most significant effect, nearly completely penetrating deep into the tumor spheroid. Control macrophages, on the other hand, exhibited only mild targeting and could seldom reach the depths of the tumor spheroid. At an E/T ratioof 10:1, CAR-HER2 macrophages demonstrated a significant cytotoxic effect on solid tumor spheroids compared to macrophages alone(Fig. 3C-D). Furthermore, we explored the infiltration and cytotoxic effects of CAR-HER2 macrophages on tumor spheroids at different time points and found that with the increase of time, the infiltration and cytotoxic effects of CAR-HER2 macrophages on the tumor spheroids gradually increased (Fig. 3E-G). Through DAPI staining and observation of macrophage GFP fluorescence, we also confirmed that CAR-HER2 macrophages successfully entered the interior of the tumor spheroids ((Fig 3H-I).

**Fig. 3.**
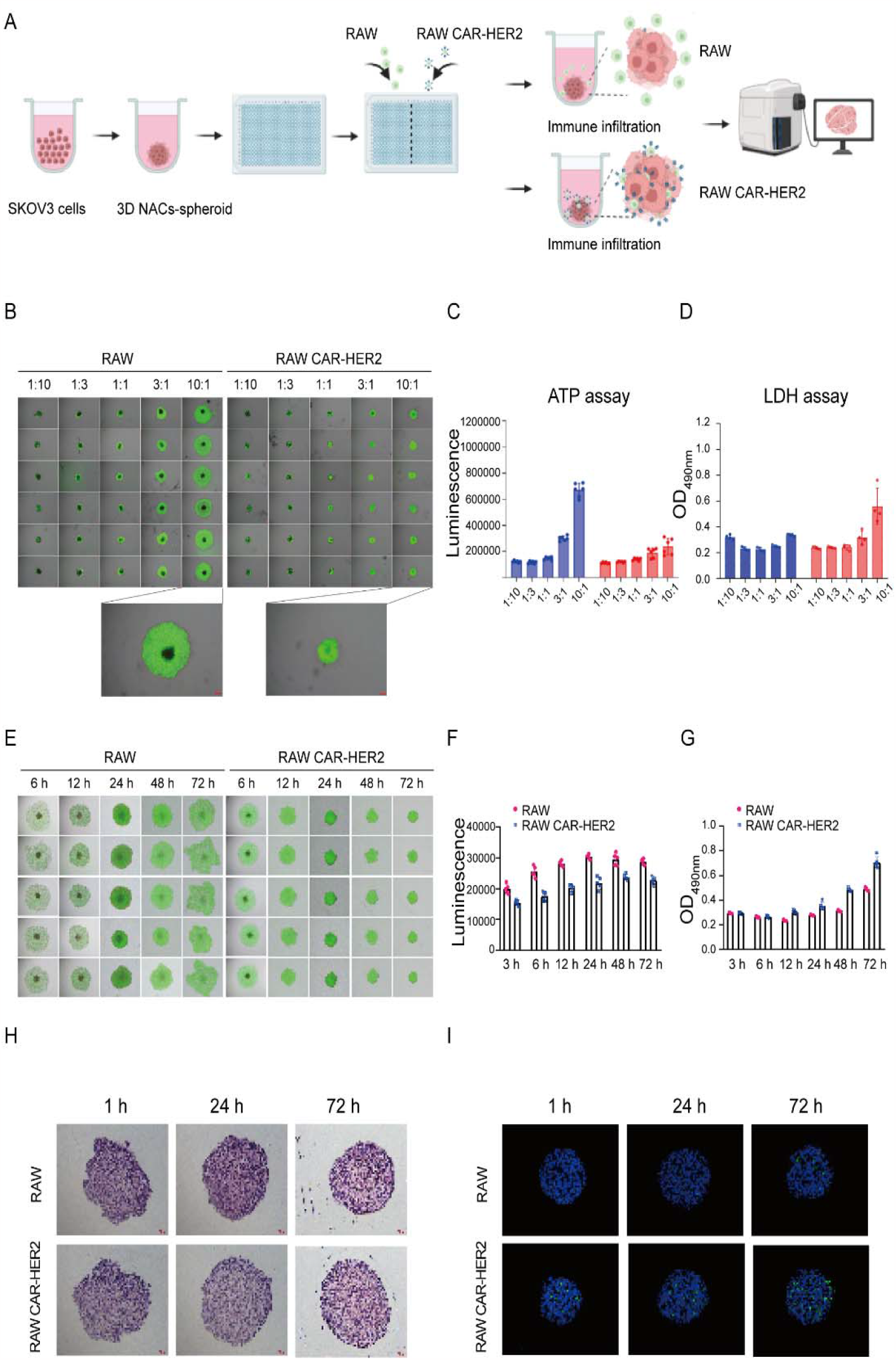
Targeting, infiltration, and cytotoxicity effects of RAW CAR-HER2 cells in 3D NACs-origami spheroid model. (3A) Schematic representation of the interaction between CAR-M and 3D NACs-origami spheroid mimicking solid tumor. (3B-D) Targeting by fluorescence microscopy using a x4 lens (3B), infiltration and cytotoxicity (3C, D) effects of CAR-M at different ratios of E/T on 3D NACs-origami spheroid. (3E-G) Targeting by fluorescence microscopy using a x4 lens, infiltration (3E), and cytotoxicity effects (3F, G) of CAR-M on 3D NACs-origami spheroid at different time points. (3H-I) H&E staining and immunofluorescent staining for DAPI at time points. Scale bar, 100 μm. For above all panels, data are represented as the mean±SD of n≥4 technical replicates experiment.

## 4. Discussion

In this study, we highlighted the effectiveness of CAR-M therapy from two aspects: a conventional 2D co-culture model and a new developed 3D tumor spheroid model based on our 3D NACs-origami assembly technique. Until now, CAR-M therapy is considered to be one of the promising treatment strategiesy in the field of solid tumor, especially in trafficking and infiltration. Likewise, our results from the conventional well/plate surface co-culture experiments showed that CAR-mediated macrophages exhibited a significant phagocytic effect compared to macrophages alone. And the killing effect varied depending on different ratios of effector to target(E/T) or effect time. In particular, we found that the most optimal efficiency was achieved when CAR-macrophages and tumor cells interacted with each other for 6 hours with an E/T ratio of 10:1 in the phagocytosis and killing assay. Furthermore, we are actively exploring ways to optimize the CAR structure through a structure-function mechanism and investigating its potential combination with other cancer therapy methods via this new 3D NACs-origami platform. This also indicates that CAR-M therapy still has huge potential for optimization.

At the same time, it was well known that periodical drug development cannot be separated from in vitro and in vivo animal experiments, and as a matter of fact, animal modeling was the most time-consuming and labor-intensive. Hence, we propose a kind of new 3D tumor micro-spheroids model explored by the NACs-origami technique. In our this study, we emphasized the feasibility of using 3D NACs-origami assembled tumor spheroids to assess the cytotoxicity and infiltration of chimeric antigen receptor macrophages. This approach serves as a link or bridge between animal and cell tests, offering the following advantages:(1)High survival rate: The 3D NACs- origami assembled tumor spheroids have a high survival rate. Our results showed that they can survive for at least 5 days, and we have successfully constructed different cell types such as hepatoma and glioma (data not shown).(2)Affinity with immune cells: The 3D NACs-origami assembled tumor spheroids have a good bio-affinity with immune cells. Our test data demonstrates that peripheral blood mononuclear cells can effectively participate in and invade the tumor spheroids. (3) Mimicking solid tumor microenvironment: The 3D NACs-origami assembled tumor spheroids also mimic a simplified version of the solid tumor microenvironment. Obviously, trafficking and infiltration barrier and tumor heterogeneity in solid tumors partially resulted in the failure of CAR-T [33].In this respect, our study had outlined that our immune cell or CAR-M could infiltrate into tumor micro-spheroid, and CAR-Ms could specifically target the solid tumor more effectively than macrophage alone. Meanwhile, CAR-M co-cultured with 3D tumor spheroids exhibited an E/T ratio- and time-dependent increase manner in targeting and infiltration efficiency, and we also found that the E/T ratio at 10:1 could reach an optimal state, which is consistent with the 2D co-culture level. However, this tumor spheroids do not fully capture the complexity of solid tumors due to the absence of heterogeneous cells. Maybe Aadding a variety of stromal cells, such as fibroblasts or, immunosuppressive tumor-associated cells and T cells may enables micro-spheroid to be closer to for mimicking solid tumor microenvironment. (4) Alternative in vitro evaluation: The 3D NACs-origami assembled tumor spheroids provides an alternative evaluation approach in vitro before preclinical animal experiments, especially in saving time and costs saving. Furthermore, combining the model with automation devices can simplify and widen its applications if some conditions permit.

In general, No matter whethereither in the traditional 2D co-culture system or the 3D tumor spheroids system, our CAR-Ms exhibited excellent targeting, infiltration, and killing capabilities against solid tumors. we also believe that this 3D NACs-based tumor spheroids system was also harnessed for other areas related to CAR-M, such as exploring combination CAR-M therapy with chemotherapy(immunomodulatory drugs)or monoclonal antibodies (e.g. anti-CD47 or Siglec proteins). Thus, the integration of these technologies together accelerates CAR-M clinical research.

## 5. Conclusion

In summary, we have established a 3D tumor spheroids system based NACs-origami assembled technique for assessing CAR-Ms in vitro. More importantly, this platform is conducive to the verification of multi-targets or multi-tumors of CAR-M and also provides reliable and favorable evidence for future clinical treatment.

## Acknowledgments

This research project was funded by the National Key Research and Development Program [2019YFA0110200, 2019YFA0110201]; and supported by Shenyang University of Chemical Technology-Roc Rock Biotechnology (Suzhou) Co., Ltd jiont reasearch grant [2021210101004604].

